# Spatial Distribution of Discarded Vehicle Tires and Their Influence on *Aedes albopictus* and *Culex quinquefasciatus* Populations, New Orleans, Louisiana

**DOI:** 10.1101/2020.02.10.942706

**Authors:** Mohamed F. Sallam, Tamer Ahmed, Cynthia Sylvain-Lear, Claudia Riegel, Imelda K. Moise

## Abstract

Discarded vehicle tires play an important role in the colonization of container mosquito populations, particularly their geographic expansion. We assessed the spatial distribution of illegally discarded tires and their response to land use-land cover (LULC), and demographic factors using geospatial analysis and generalized regression. Multiple stepwise regressions were used to evaluate the response of the Container Index (CI) of colonized *Aedes albopictus* (Skuse), and *Culex quinquefasciatus* Say to macro- and microhabitats variables. The illegally discarded tires were distributed over 11 planning districts with clustering distribution for tires frequency and overdispersed distribution for tires number. Out of 1,137 (∼37.08%) water-holding tires, 598 (∼52.64%) tires at 65 (∼38.46%) sites were positive for colonized mosquito populations. A total of 13 mosquito species were identified, with the highest CI of *Ae. albopictus* (44.19%) and *Cx. quinquefasciatus* (22.18%). *Aedes albopictus* colonized all 65 sample sites and *Cx. quinquefasciatus* found at 32 sites. The Container Index (CI) of colonized mosquito was clustered in seven planning districts for *Ae. albopictus* and five planning districts for *Cx. quinquefasciatus*. Microhabitat (muddy water) rather than macrohabitats variables predicted both species’ colonization, especially *Ae. albopictus*. The contribution of macro- and microhabitat characteristics in predicting colonized mosquito in water-holding tires was discussed.

## Introduction

Discarded vehicle tires play an important role in the proliferation of mosquito colonization and breeding (Shannon 1931), particularly the introduction and geographic expansion of mosquito species. The most striking example of this is the introduction of the Asian tiger mosquito, *Aedes albopictus* (Skuse) (Hawley et al. 1987) and *Ae. (Ae) japonicus* (Theobald) (Peyton et al. 1999) to the United States (US) from Asia and South Pacific. Tires are also a habitat for other mosquito species in the continental US (e.g., *Culex (Cx.) restuans, Cx. pipiens, Cx. territans, Cx. Salinarius and Cx. quinquefasciatus*) (Yee 2008). The introduction of *Ae. albopictus* to North America after World War II set the stage for the transmission of arboviruses, causing both wildlife and human disease outbreaks such as dengue, chikungunya (Paupy et al. 2009), and La Crosse encephalitis (Gerhardt et al. 2001).

Of particular concern is the durability of tires as water-holding habitats as containers for mosquitos that vector diseases. Because of their specific shape and impermeable nature, they can produce adult mosquitoes over longer periods than natural containers (e.g., leaf axils, tree holes) (Sofi 2018) or even other artificial containers such as plastic trash and food containers (Dowling et al. 2013). This has been relevant to public health because tires are most likely to be discarded near human populations and hence pose health risks (Yee 2008, Villena et al. 2017). Studies over the past two decades have provided important information on the link between the location of tires and female oviposition behavior (Kling et al. 2007, Day 2016, Allgood and Yee 2017), but few have investigated variation in the density and distributions of larvae in discarded tires (Lampman et al. 1997, Qualls and Mullen 2006).

While other studies highlighted the spatial distribution of mosquito populations in tire systems and their response to environmental variables in the U.S (Juliano and Lounibos 2005, Qualls and Mullen 2006, Bartlett-Healy et al. 2012, Yee et al. 2012) and Latin America (Manrique-Saide et al. 2008, Troyo et al. 2008, Mendonça et al. 2011, García-Rejón et al. 2012, Baak-Baak et al. 2014), they did not give detailed information on i) the frequency and density of discarded vehicle tires, ii) the general effect of the interaction between landscape and demographic subsystems on discarded tires and mosquito populations, and iii) the interplay between macro- and microhabitat characteristics and their impact on mosquito populations.

In this study, we evaluated the geospatial distribution patterns of frequency and number of discarded vehicle tires, and tires response to land use-land cover (LULC), and demographic predictors. Additionally, we assessed the response of the container index of colonized *Ae. albopictus* and *Cx. quinquefasciatus* populations to their predicting factors within macro- and microhabitat characteristics. Finally, we evaluated the interaction between macro- and microhabitat characteristics affecting the colonization of these two mosquito populations.

## Materials and Methods

### Study Area

The city of New Orleans Louisiana (NOLA) lies on the Mississippi River, near the Gulf of Mexico with a total land area of 438.80 km^2^. In 2017, NOLA was inhabited by almost 343,829 people, with an average density of 892.2/km^2^. NOLA is sub-tropical with an annual high temperature of 25^°^C, an annual low of 16.8^°^C, and average annual precipitation of 162.3 cm. The average highest precipitation occurs in July (17.9 cm).

### Discarded Vehicle Tire Surveillance

We obtained 311 records of illegal discarded tire sites from the City of New Orleans through crowdsourcing data from local government and non-emergency information. The City of New Orleans Department of Sanitation reported and confirmed the calls regarding illegally discarded tires reported to the city’s 311-call center or via email by residents, visitors, or businesses (www.nola.gov/311).

A purposive tire sampling based on 311 records was conducted for confirmation. Additionally, random surveillance was conducted within the reported housing blocks and neighborhood of the reported sites: i) to explore other dumping sites that have not been reported, and ii) to represent different landscape characteristics of dumping sites. The random surveillance was conducted within pre-defined sampling grids using the sampling design tool in ArcGIS ver. 10.1 (ESRI 2011) (Figure 1). Data records for discarded vehicle tires were compiled from both the 311-call database and active random surveillance during January 2015-January 2018. During this study, we did not account for legally retained or illegally dumped tires in areas not accessible such as fenced private premises.

**Figure 1.**
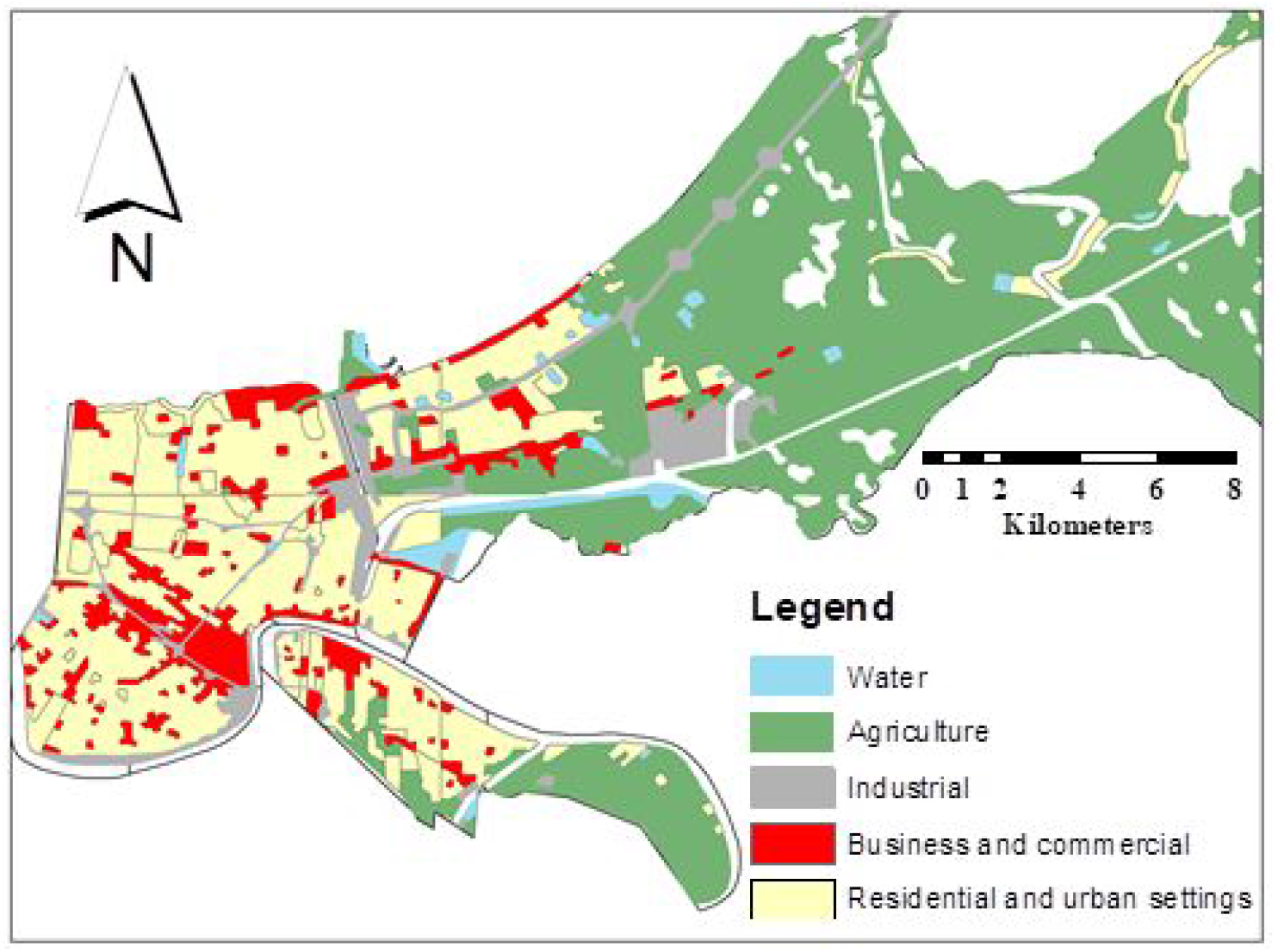
Map of the city of New Orleans showing different land use-land cover and random sampling points generated for surveillance.

### Mosquito Sampling

All water-holding tires were sampled for mosquito colonization and container index (CI) estimates from November 2016 to January 2018. During surveillance, three types of tires were sampled: A) light truck, 8-ply non-radial, 70 cm diam × 17 cm wide; B) radial automobile, 56 × 17.5 cm, and C) tractor, 128 × 70 cm. Physical characters of water, such as pH, temperature, and water quality, were collected before sampling the colonized mosquitos. Tire water was mixed before sampling to obtain bottom-dwelling mosquito immatures. A destructive sampling method was used to collect all biological materials from water-holding tires. All water contents from the tires were strained and transferred into Nasco Whirl-Pak bags to the New Orleans Mosquito, Termite Control Board (NOMTCB) laboratory for further sample assortment, and identification. The number of larvae and pupae was quantified at our facility. Only fourth larval instars were used for the identification of species. Early instars that were not readily identifiable were reared in plastic mosquito breeders at 27^°^C±2 and 80±5 RH% to the fourth instar before being identified. Pupae were similarly maintained and identified as adults. Larval and adult stages were identified by species level using the taxonomic keys of Darsie and Ward (Darsie and Ward 2005). All possible predators were quantified and recorded for each sampled tire.

### Microhabitat Data Variables

Microhabitats variables were represented as seven biophysical factors: size of tires (small, large, mixed), tire status (damaged, intact), water temperature, pH value, water quality (muddy, turbid, clear), predators (midges, water fleas, midges/water fleas), and CI of colonized mosquito species (Table 1).

**Table 1.**
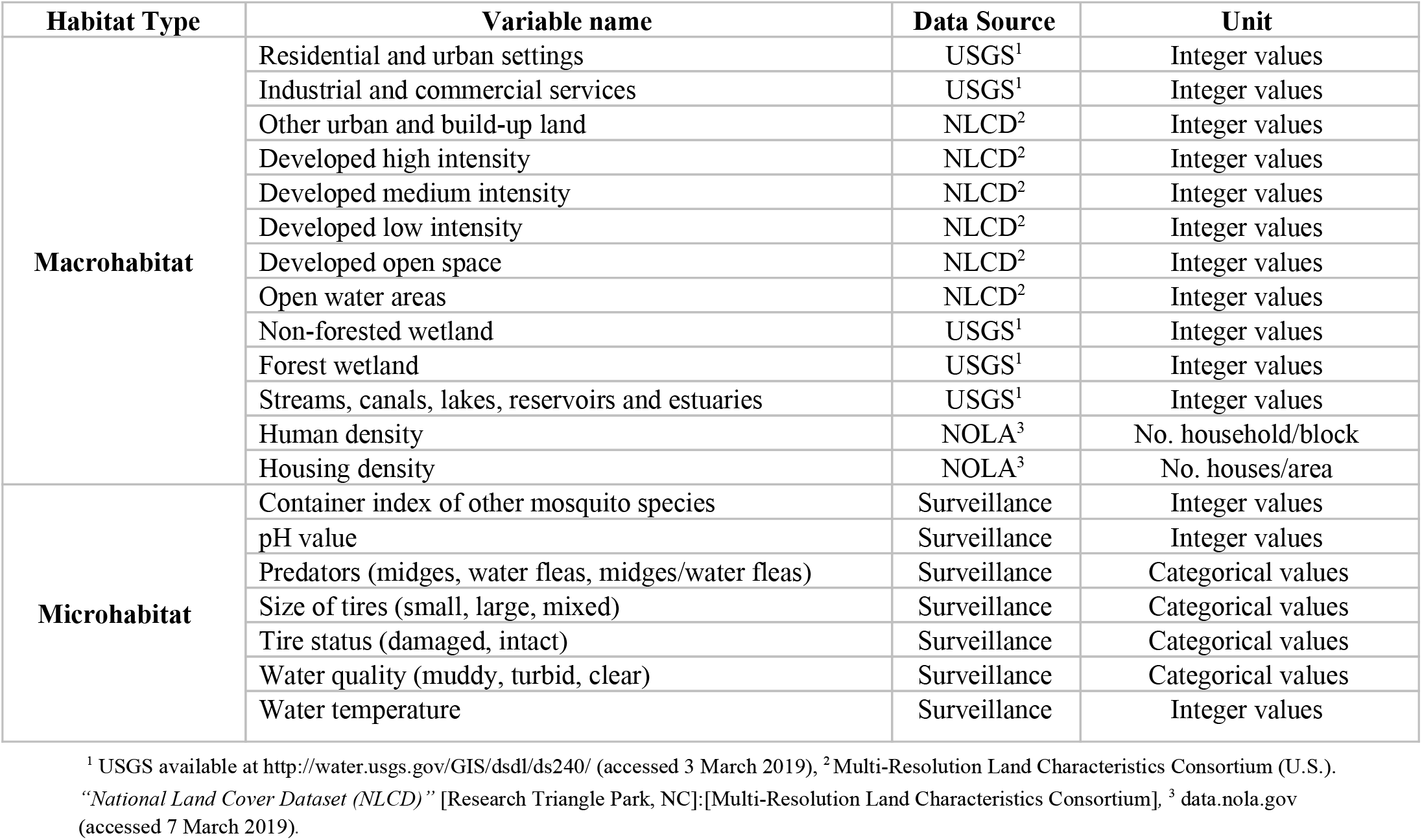
Proposed macro- and microhabitat variables used in regression models.

| Habitat Type | Variable name | Data Source | Unit |
| --- | --- | --- | --- |
| Macrohabitat | Residential and urban settings | USGS<sup>1</sup> | Integer values |
|  | Industrial and commercial services | USGS<sup>1</sup> | Integer values |
|  | Other urban and build-up land | NLCD<sup>2</sup> | Integer values |
|  | Developed high intensity | NLCD<sup>2</sup> | Integer values |
|  | Developed medium intensity | NLCD<sup>2</sup> | Integer values |
|  | Developed low intensity | NLCD<sup>2</sup> | Integer values |
|  | Developed open space | NLCD<sup>2</sup> | Integer values |
|  | Open water areas | NLCD<sup>2</sup> | Integer values |
|  | Non-forested wetland | USGS<sup>1</sup> | Integer values |
|  | Forest wetland | USGS<sup>1</sup> | Integer values |
|  | Streams, canals, lakes, reservoirs and estuaries | USGS<sup>1</sup> | Integer values |
|  | Human density | NOLA<sup>3</sup> | No. household/block |
|  | Housing density | NOLA<sup>3</sup> | No. houses/area |
| Microhabitat | Container index of other mosquito species | Surveillance | Integer values |
|  | pH value | Surveillance | Integer values |
|  | Predators (midges, water fleas, midges/water fleas) | Surveillance | Categorical values |
|  | Size of tires (small, large, mixed) | Surveillance | Categorical values |
|  | Tire status (damaged, intact) | Surveillance | Categorical values |
|  | Water quality (muddy, turbid, clear) | Surveillance | Categorical values |
|  | Water temperature | Surveillance | Integer values |

<sup>1</sup> USGS available at http://water.usgs.gov/GIS/dsdl/ds240/ (accessed 3 March 2019), <sup>2</sup> Multi-Resolution Land Characteristics Consortium (U.S.). "National Land Cover Dataset (NLCD)" [Research Triangle Park, NC]:[Multi-Resolution Land Characteristics Consortium], <sup>3</sup> data.nola.gov (accessed 7 March 2019).

### Macrohabitat Data Variables

#### Land Use-Land Cover (LULC) and Demographic Data

Because adult female mosquitoes require a blood meal for egg development (Zhu et al. 2014), we derived the recently updated (2007) LULC from the US Geological Survey (USGS) (Burkett-Cadena et al. 2010) to delineate land use and land cover patterns within the New Orleans metropolitan area, to identify areas with a higher likelihood of finding a blood meal. Urban areas were categorized into three classes to represent the degree of urbanization: (1) residential and urban, in which housing predominates in two different forms; (2) industrial and commercial services, which represents industrial settings and fewer housing structures; and (3) other urban and built-up land with the least housing and human populations. We then used the National Land Cover Database 2011 (NLCD) to define five major classes of LULC to represent environmental predictors linked to high vector densities (Ruiz et al. 2004). These five major classes are: 1) developed high intensity, 2) developed medium intensity, 3) developed low intensity, 4) developed open space, and 5) open water areas.

Further, because vegetated vacant lots at sample sites may represent resting places and sugar meal sources for adult mosquitoes (Sallam et al. 2016) and possible disposal sites, two major classes of vegetation types were extracted from the USGS (Lewis et al. 2017). The two extracted vegetation types include non-forested wetlands and forest wetlands. Further, we included streams, canals, lakes, reservoirs, and estuaries of different sizes (>1 and ≤1 km) to represent permanent water bodies as possible illegal discarded tire sites (Figure 1). Demographic data (human population and housing density) at the block level was downloaded from the US Census American Community Survey (ACS) for 2012-2016.

### Data Analysis

#### Geospatial Distribution Analysis

We were specifically interested in evaluating the geospatial distribution of frequency (number of dumping incidences per site) and the number of illegally discarded tires. Additionally, we assessed the CI of two dominant mosquito populations in the tire systems. First, we measured the geographic distribution of illegally discarded tires within areas with an elevated risk of having a high frequency of discarded tires and reported the number using the standard deviational ellipse (SDE) statistical tools in ArcGIS. Second, we used Global Moran’s I (Anselin and Getis 1992), a correlation coefficient that measures the overall spatial autocorrelation of locations (e.g., illegal tire sites) and feature values (e.g., frequency and reported the number of tires) concurrently. Clusters of sites with a high frequency and density of illegally discarded tires were considered “high-risk areas” whereas clusters of sites with low frequency and density of illegally discarded tires (Low-Low) were considered “low-risk sites.” The value of Moran’s I is measured between −1 and 1, with the ‘Z’ score value calculated to test whether the observed clustering or dispersal is significant. Therefore, when the Z score indicates statistical significance, a positive Moran’s I indicates a tendency toward clustering while a negative value indicates a tendency toward dispersion. When the Z score value is not significantly different from zero, it is assumed that there is no spatial autocorrelation, and the pattern does not appear to be significantly different from a random distribution.

Similarly, the spatial distribution and Global Moran’s I index for the CI of *Ae. albopictus* and *Cx. quinquefasciatus* populations were evaluated. The clustering hypothesis (H_1_) was tested for the frequency and density of discarded vehicle tires and the colonized mosquito populations. The null hypothesis (Ho) was that there is no spatial clustering of frequency and density of discarded vehicle tires or the CI of colonized mosquito populations in the city of New Orleans.

### Statistical Analysis

To characterize the influence of the 13 LULC and demographic predictors that predict discarded tire frequency and number, two generalized regression models (GRM) with lasso tool were carried out to delineate their response to predicting factors from the sampling sites.

On the other hand, multiple stepwise linear regressions were conducted to evaluate: (1) the influence of macrohabitat characteristics on the CI of *Ae. albopictus* and *Cx. quinquefasciatus* population, (2) the influence of microhabitat characteristics on the CI of *Ae. albopictus* and *Cx. quinquefasciatus* colonization, and (3) the interaction between macro- and microhabitat predictors and their influence on the CI of *Ae. albopictus* and *Cx. quinquefasciatus* in water-containing tires. For colonized mosquito populations, macrohabitats were denoted as a total of 13 LULC and demographic data layers, and microhabitats variables were represented as seven biophysical factors (Table 1).

The log10(n+1) formula was used to transform the number of tires and area percentages of LULC to maintain the normality of their statistical distribution. All statistical analyses were conducted in SAS 14.0 (SAS 2013) using the JMP pro statistical package. Significant predictors and best prediction models were selected based on minimum corrected Akaike Information Criterion (AICc) and maximum *r*^2^ values, with *p* < 0.05. All variables were standardized for the average and standard deviation become 1 and 0.

## Results

### Dumping Tires and Mosquito Sampling

Four hundred and five sites with 6,491 discarded tires were surveyed during the current study representing 13 New Orleans planning districts. A total of 236 (∼58.27%) sites with 3,427 (∼52.80%) tires were reported by the 311-phone service. Additionally, 3,064 (∼47.20%) discarded tires were actively surveyed for mosquito immature colonization from 169 sites (∼41.73%). The dumping incidences of tires reported from all sites were either one or two times, representing 64 and 341 sites, respectively.

Tire discarding sites reported by 311 services could not be surveyed for mosquito immatures because the department of sanitation of the city of New Orleans removed tires at these sites. Of the 3,064 surveyed tires, 1,136 (∼37.08%) were found to contain water, and 598 (∼52.64%) of water containing tires were positive for mosquito immatures colonization.

We observed positive mosquito colonization at 65 (∼38.46%) sampling sites with 2,900 mosquito immatures (larvae/pupae) in sampled tires. These tires provided suitable colonization habitats for 13 mosquito species with a new locality record of *Aedes japonicus*^35^ (Table 2). Colonized *Ae. Albopictus* populations were the highest (CI = ∼44.19), followed by *Cx. quinquefasciatus* (CI = ∼22.18). *Aedes albopictus* was recorded from 502 tires at 65 sites. Whereas, *Cx. quinquefasciatus* was collected from 252 tires at 32 sites. The dominance *of Ae. albopictus* population also was demonstrated on spatial scale, as it was collected from seven districts (∼54%). In the meantime, *Cx. quinquefasciatus* was reported in five districts (∼39%). Both species co-existed in 21 sampling sites distributed in four planning districts: Bywater, Gentilly, Lower Ninth Ward, and New Orleans East.

**Table 2.**
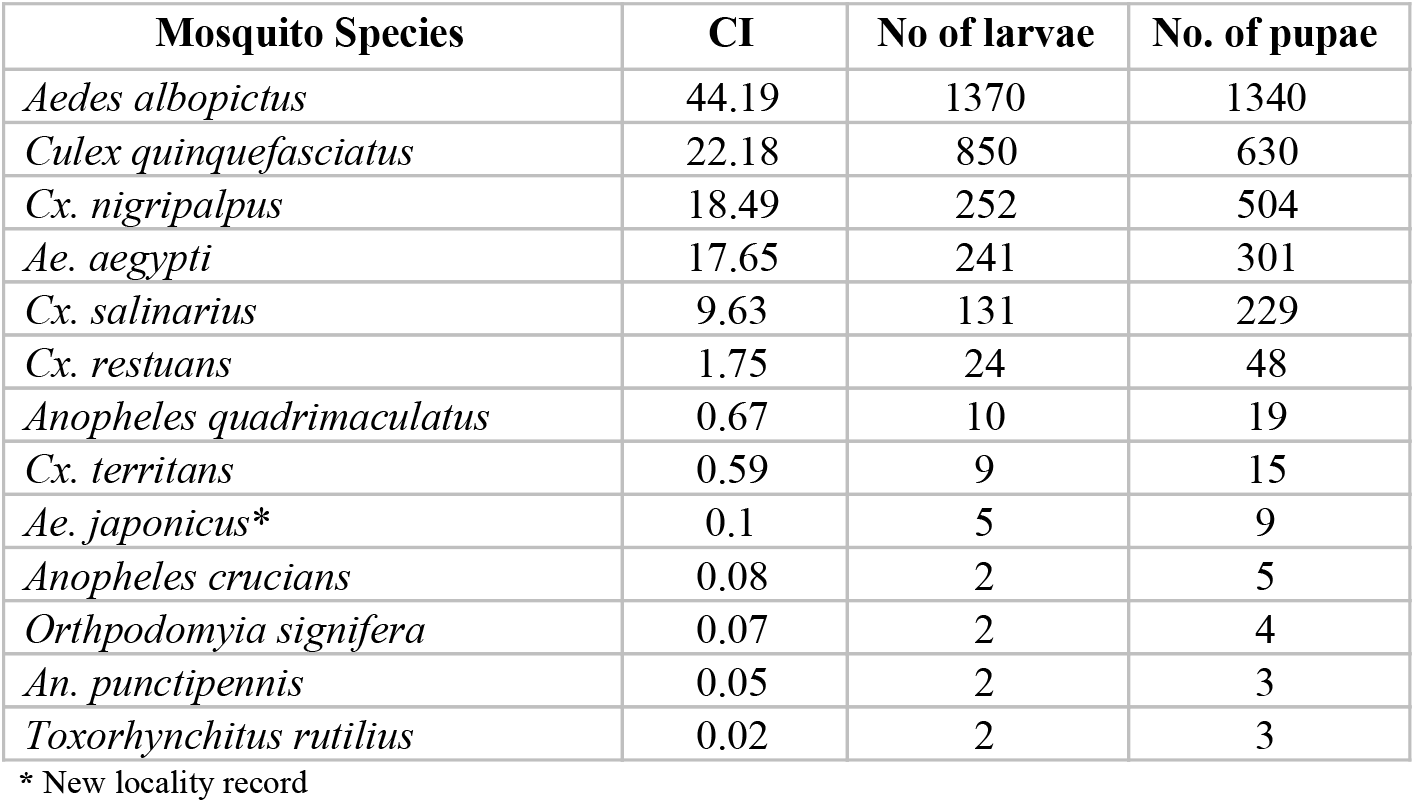
Container Index (CI), number of larvae and pupae sampled from discarded tires in the City of New Orleans, LA

| Mosquito Species | CI | No of larvae | No. of pupae |
| --- | --- | --- | --- |
| *Aedes albopictus* | 44.19 | 1370 | 1340 |
| *Culex quinquefasciatus* | 22.18 | 850 | 630 |
| *Cx. nigripalpus* | 18.49 | 252 | 504 |
| *Ae. aegypti* | 17.65 | 241 | 301 |
| *Cx. salinarius* | 9.63 | 131 | 229 |
| *Cx. restuans* | 1.75 | 24 | 48 |
| *Anopheles quadrimaculatus* | 0.67 | 10 | 19 |
| *Cx. territans* | 0.59 | 9 | 15 |
| *Ae. japonicus* * | 0.1 | 5 | 9 |
| *Anopheles crucians* | 0.08 | 2 | 5 |
| *Orthopodomyia signifera* | 0.07 | 2 | 4 |
| *An. punctipennis* | 0.05 | 2 | 3 |
| *Toxorhynchitis rutilius* | 0.02 | 2 | 3 |

\* New locality record

### Geospatial Distribution Analysis

We used the Moran’s I to measure global clustering patterns of illegal discarded tire frequency. We found spatial clustering of dumping frequency (Z-score = 1.76, *p*<0.10). Most of these discarded tires were in areas of the 13 New Orleans planning districts (Figure 2). In contrast, the number of discarded tires showed a random spatial distribution (Z-score value = −0.17, *p*>0.05). However, the concentration centers of illegally discarded tire frequency and reported numbers were in Mid-city. We also found that certain areas of New Orleans have an elevated risk for discarded tires in both frequency and numbers. This area covers a total area of 24.8 Km^2^ (5.64%) and 23.3 Km^2^ (5.31%) of the total land area of New Orleans and is mostly located in 11 planning districts.

**Figure 2.**
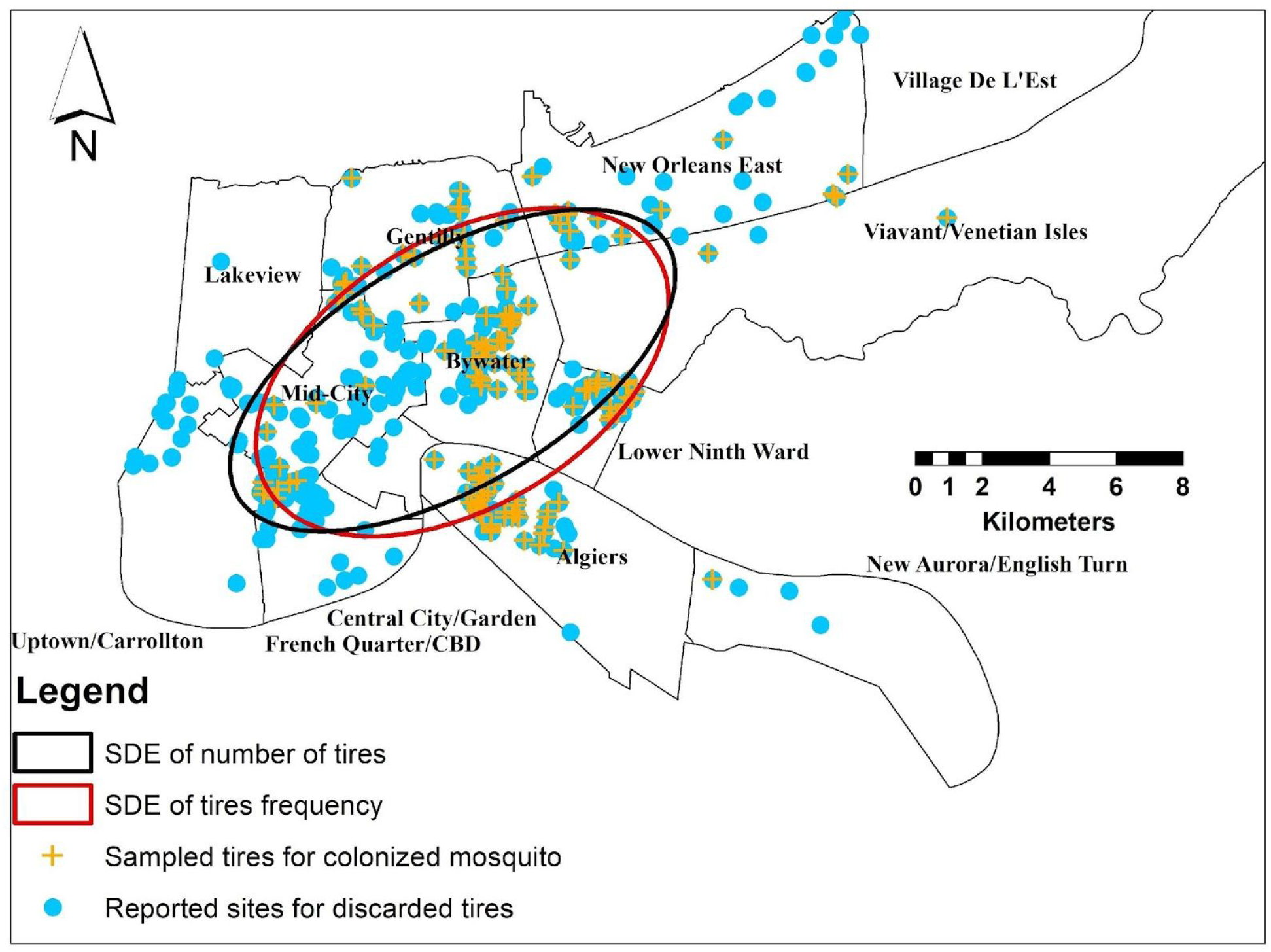
Standard Deviational Ellipse (SDE) for dumping frequency and number of tires.

Similarly, spatial autocorrelation and geographic distribution patterns for the CI of colonized *Ae. albopictus* and *Cx. quinquefasciatus* populations were evaluated to test the clustering hypothesis (H_1_). Only the CI of *Ae. albopictus* showed the significant clustered distribution and the center of concentration was found to be in Bywater.

Meanwhile, the CI of *Cx. quinquefasciatus* demonstrated a random distribution pattern, and the concentration center of its distribution also was found to be in Bywater. The distance between mean centers of concentration for both species was 0.47 Km.

The clustered and random distribution patterns were confirmed by the significant positive Z-score values (*Ae. albopictus*= 3.61, *Cx. quinquefasciatus*= 1.62, *p* <0.05). Accordingly, the clustering hypothesis (H_1_,) for the CI of *Ae. albopictus* colonized in discarded tires was accepted. The geospatial distribution demonstrated by SDE polygons showed areas under high risk of increased CI of colonized *Ae. albopictus* and *Cx. quinquefasciatus* covered 55.19 Km^2^ (∼12.58%) and 41.88 Km^2^ (∼9.54%), respectively, of the total area of the city. Areas under risk of both mosquito populations that colonized discarded tires were found to represent 39.95 Km^2^ (9.10%) of the studied sites. These areas were found to represent seven planning districts: Bywater, Gentilly, Lower Ninth Ward, Mid-City, New Orleans East, Algiers, and Venetian Isles (Figures 3 and 4). The SDEs of both mosquito populations were found to overlap in an area of 36.06 Km2 (∼8.22%). This finding reflects the co-existence of both species within the same sampling sites and tire habitats.

**Figure 3.**
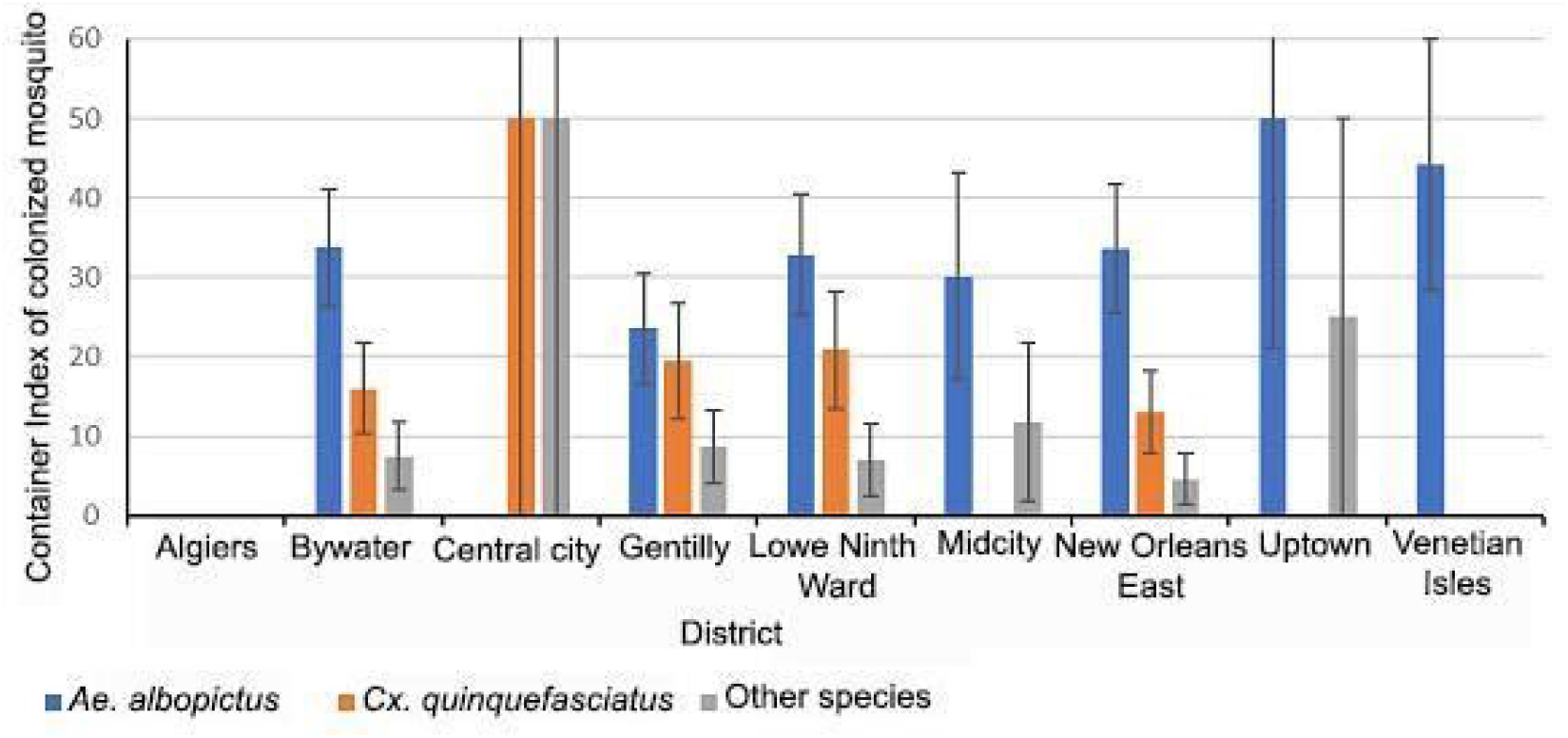
Spatial distribution of colonized mosquito populations in the city of New Orleans.

**Figure 4.**
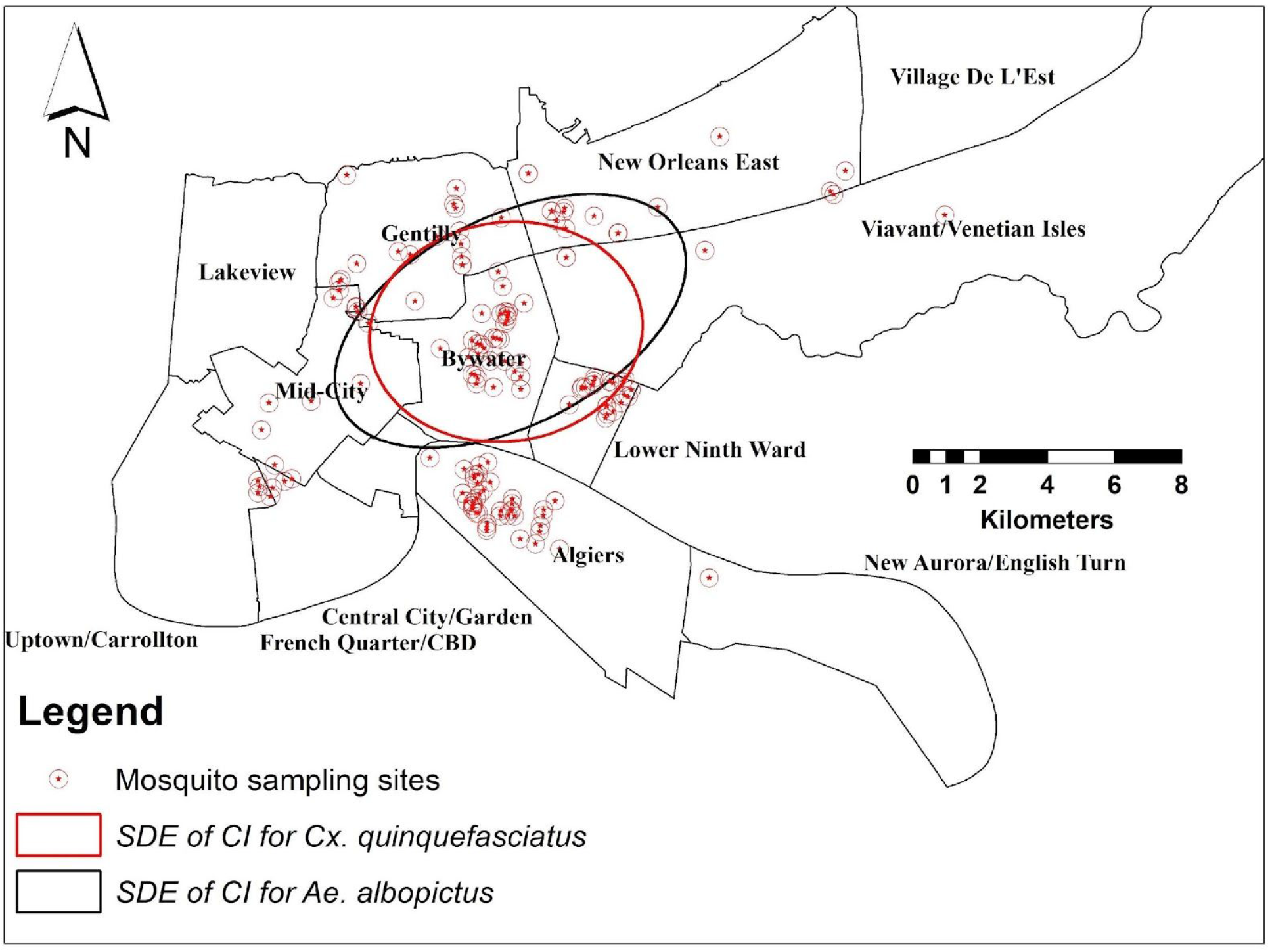
Standard Deviational Ellipse (SDE) of Container Index of colonized mosquito populations in the city of New Orleans.

### Statistical Regression Models

#### Spatial Characteristics of Illegal Discarded Tires

Two factors were significantly associated with illegally discarded tire frequency (r^2^=4.58%, AICc=321.63, *p*<0.05). Industrial areas were positively associated with discarded tire frequency (Table 3). None of LULC variables showed any significant association with discarded tire frequency. Meanwhile, five variables were significantly associated with the increased number of dumping tires (r^2^=3.52%, AICc=1237.24, *p*<0.05). All variables showed a negative association with an increased number of tires dumping with the exception of industrial areas (Table 3).

**Table 3.** The response of frequency and number of increased dumping tires to landscape and demographic variables.

| Model Type | Influential Factor | $\beta$ | Wald $\chi^2$ |
| --- | --- | --- | --- |
| Frequency of dumping tires | Industrial areas | 1.72E-7 | 6.32** |
|  | Average housing density | -0.01 | 6.45** |
| Number of dumping tires | Other urban built-up areas | -7.4E-6 | 22.15* |
|  | Average housing density | -0.06 | 7.47* |
|  | Industrial areas | 6.66E-7 | 5.11** |
|  | Residential and urban settings | -9.42E-7 | 5.01** |
|  | Non-forested areas | -2.62E-7 | 3.91** |

\* $p < 0.01$ , \*\* $p < 0.05$

#### Spatial Characteristics of Mosquito Populations

The scale of regression models, whether macro-or microhabitats, influenced the number and type of predicting variables for the colonization of each mosquito species. For *Ae. albopictus*, two macrohabitat variables have a significant maximum influence in predicting the CI. These variables were developed medium intensity (β *=* 7.16E-6, *r*^2^=5.23%, AICc=1674, *p*<0.05), and developed low intensity (β= −1.75E-6, *r*^2^=6.64%, AICc=1673, *p*<0.05) as indicated by *r*^2^ and AICc values. Developed low-intensity areas were found to be negatively associated with the increased CI of colonized *Ae. albopictus* population (Table 4).

**Table 4.**
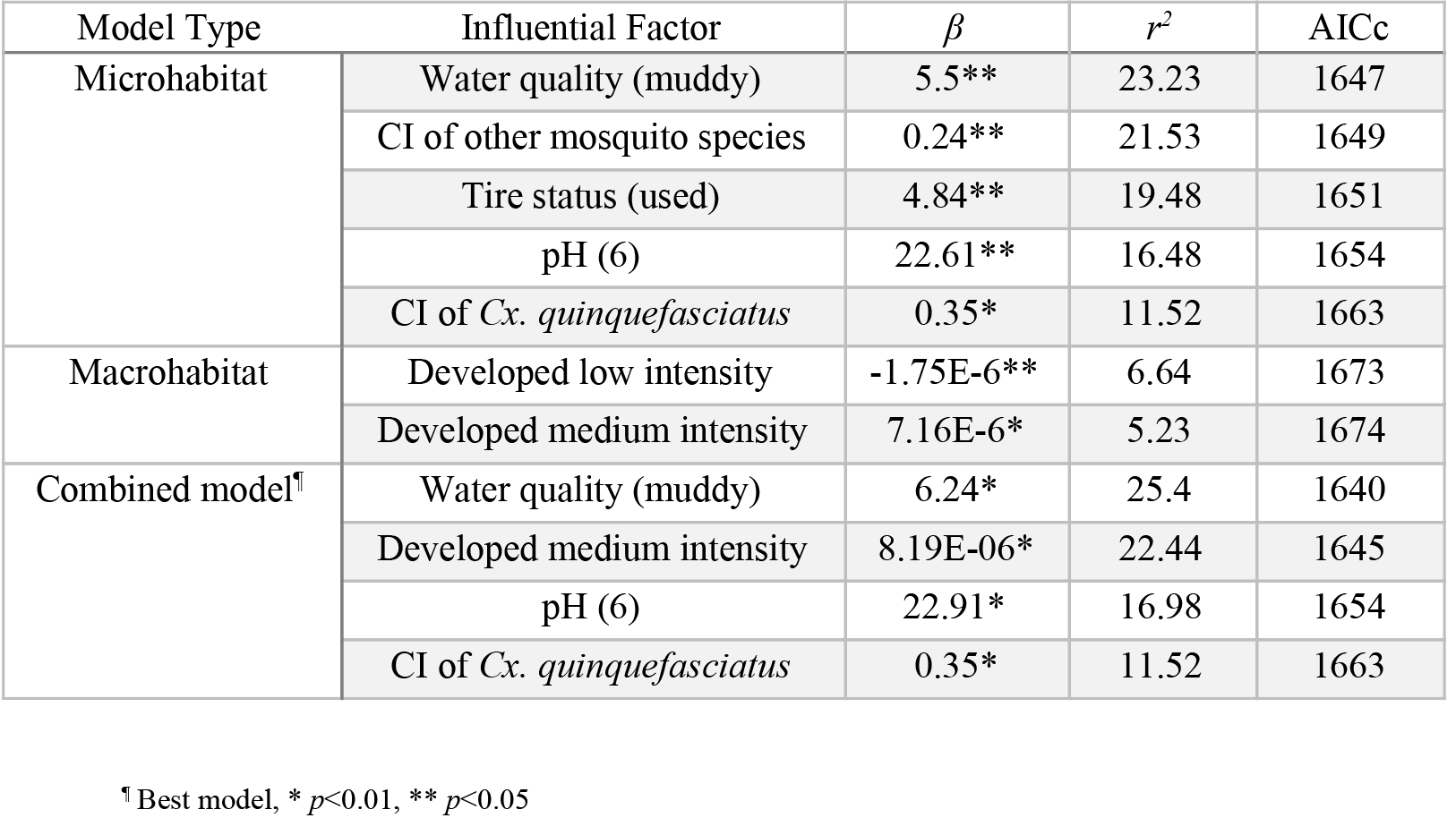
The response of colonized *Ae. albopictus* population to micro- and macrohabitat variables.

| Model Type | Influential Factor | $\beta$ | $r^2$ | AICc |
| --- | --- | --- | --- | --- |
| Microhabitat | Water quality (muddy) | 5.5** | 23.23 | 1647 |
|  | CI of other mosquito species | 0.24** | 21.53 | 1649 |
|  | Tire status (used) | 4.84** | 19.48 | 1651 |
|  | pH (6) | 22.61** | 16.48 | 1654 |
|  | CI of *Cx. quinquefasciatus* | 0.35* | 11.52 | 1663 |
| Macrohabitat | Developed low intensity | -1.75E-6** | 6.64 | 1673 |
|  | Developed medium intensity | 7.16E-6* | 5.23 | 1674 |
| Combined model<sup>¶</sup> | Water quality (muddy) | 6.24* | 25.4 | 1640 |
|  | Developed medium intensity | 8.19E-06* | 22.44 | 1645 |
|  | pH (6) | 22.91* | 16.98 | 1654 |
|  | CI of *Cx. quinquefasciatus* | 0.35* | 11.52 | 1663 |

<sup>¶</sup> Best model, \* $p < 0.01$ , \*\* $p < 0.05$

On the other hand, the microhabitat variables demonstrated an increased prediction effect on the CI of colonized *Ae. albopictus* (*r*^2^= 23.23%, AICc=1647, *p*<0.05). Five predicting variables were found to increase colonized *Ae. albopictus* populations (Table 4). The water quality demonstrated the maximum prediction power with the highest *r*^2^ and minimum AICc values (β*=* 5.5, *r*^2^=23.23%, AICc=1647, *p*<0.05). Additionally, the presence of other colonized mosquito populations in water-containing tires used tire status and water alkaline pH value shared reduced prediction gain in increasing CI of colonized *Ae. albopictus*.

The combined regression model approach highlighted how the interaction between macro- and microhabitat variables could change the number and type of influential factors that affect mosquito density. The combined regression model showed the best prediction power in characterizing suitable habitats for the colonization of *Ae. albopictus* (*r*^2^= 25.40%, AICc=1640, *p*<0.01). The combined model demonstrated the contribution of three microhabitats and one macrohabitat variables in affecting the CI (Table 4). Accordingly, the combined regression approach was selected as the best prediction model because of the recorded minimum AICc value for the most predictive variable.

For *Cx. quinquefasciatus*, the developed medium intensity areas were the only macrohabitat variable that significantly predict increased CI values (β= 1.69, *r*^2^=6.0%, AICc=1673, *p*<0.05). Four microhabitat variables were found to be the most influential factors in predicting increased CI values (*r*^2^=18.68%, AICc=1567, *p*<0.05). Muddy and clear water quality demonstrated the highest prediction gain in affecting the colonization of *Cx. quinquefasciatus* (β= 6.46, *r*^2^=18.68%, AICc=1567, *p*<0.05). Additionally, intact tire status (undamaged), large tire size (tractor, 128 × 70 cm), and CI values of colonized *Ae. albopictus* were positively correlated with increased CI values of *Cx. quinquefasciatus* (Table 5). The combined regression model showed that the four microhabitat variables superseded the developed medium intensity areas in predicting increased CI values. The combined regression approach was selected as the best prediction model because of the recorded minimum AICc value.

**Table 5.** The response of colonized *Cx. quinquefasciatus* population to micro- and macrohabitat variables.

| Model Type | Influential Factor | $\beta$ | $r^2$ | AICc |
| --- | --- | --- | --- | --- |
| Microhabitat | Water quality (muddy and clear) | 6.46** | 18.67 | 1567 |
|  | Tire status (intact) | 5.42** | 17.01 | 1568 |
|  | Tire size (large) | 6.68** | 14.21 | 1572 |
|  | CI of *Ae. albopictus* | 0.25* | 11.52 | 1575 |
| Macrohabitat | Developed medium intensity | 1.69** | 6.00 | 1673 |
| Combined model<sup>†</sup> | Water quality (muddy and clear) | 4.42** | 18.67 | 1566 |
|  | Tire status (intact) | 5.14** | 17.01 | 1568 |
|  | Tire size (large) | 7.35** | 14.21 | 1572 |
|  | CI of *Ae. albopictus* | 0.26* | 11.52 | 1575 |

<sup>†</sup> Best model, \* $p < 0.01$ , \*\* $p < 0.05$

## Discussion

To our knowledge, this study is the first in New Orleans to investigate geographic distribution patterns, habitat characteristics and ecological suitability of illegally discarded vehicle tires and their associated mosquito populations. Previous work in New Orleans highlighted natural *Ae. albopictus* population in tires (Marten 1990b, a, Comiskey et al. 1999) and container-breeding *Ae. aegypti* (Focks et al. 1981). Focks et al. (1981) highlighted the significance of tires as compared to other available water-holding containers as potential breeding sites for *Ae. aegypti* and *Cx. quinquefasciatus*. Other local reports conducted by the City of New Orleans and department of health concluded that tires are strong breeding sites for mosquitoes as they are easily filled with water and collect leaf litter, providing a prime spot for mosquito vectors that can transmit West Nile, Zika and dengue (NOLA 2019). However, the previous studies either focused on biological control or the biology of these mosquito populations in tire systems. Our preliminary findings characterized areas under risk of increased frequency and number of discarded tires and two mosquito vectors populations. We investigated the number and type of influential factors that predict the container index (CI) of colonized *Ae. albopictus* and *Cx. quinquefasciatus* populations within their macro- and microhabitats. We also highlighted the significance of evaluating the interaction between macro- and microhabitat variables in predicting increased CI values of the colonized mosquito populations.

The conflicting findings between the clustered patterns of the dumping frequency and their number are possible because of the impact of landscape and demographic factors such as housing density or vegetation. Adding the number of tires to the algorithm of spatial autocorrelation reflects the magnitude of influence of certain landscape and demographic factors. For example, average housing density, other built-up urban and residential/urban were negatively associated with increased dumping frequency and number of discarded tires. A possible explanation for this is that highly populated areas were either not suitable as dumping sites or people have the tendency to discard tires in areas with low population density. This finding would improve our understanding of their influence in predicting the likelihood of an increased number of dumped tires. This would also shed light on the significant negative association of other urban built-up areas and average housing density with the dumping problem in certain areas. We recommend prioritizing neighborhoods with similar LULC types for surveillance and control efforts. The current study found that industrial areas were positively associated with both discarded tire frequency and number. No correlations were found between the frequency or number of dumped tires and population density. Additionally, the negative associations between the number of dumped tires and housing density, or residential and urban settings reflect that the number of dumping reports were not correlated with populated areas. Previous studies in other areas showed that the dumping frequency and number of tires were positively associated with grown vegetation in vacant lots and/or community perceptions, as people would store a variety of containers in their yard (Andreadis 1988, Mazine et al. 1996). For example, discarded tire frequency and density increased in neighborhoods with lower housing density. This finding is confirmed by the significant negative association between developed medium areas with increased discarded tire frequency and number. These patterns of discarded tire frequency and number in New Orleans have ecological consequences that affect both ecosystem services and human health (Lewis et al. 2017). However, further studies are needed to include the influence of other socioeconomic factors such as community perception, income rates, and education level. This not only will help in highlighting high risk areas of dumping frequency but also will shed the light on community engagement in the proliferation of illegally discarded tires.

Out of 65 positive mosquito sites, both *Ae. albopictus* and *Cx. quinquefasciatus* co-existed in 21 sampling sites representing 214 tires (35.79%) from four planning districts: Bywater, Gentilly, Lower Ninth Ward, and New Orleans East. The clustering pattern of CI for *Ae. albopictus* demonstrates their association with certain biophysical characters that predict their distribution. Meanwhile, the random distribution pattern of *Cx. quiqnuefasciatus* generated by standard deviational ellipsoid may indicate their response to microhabitat characteristics or the contribution of other potential habitats that predict their colonization, which should be addressed in a separate investigation.

The consistent interactions between macro- and microhabitat variables were demonstrated by the variation in the number and type of variables that predict the CI of mosquito populations. For example, our regression models showed that the distribution range of *Ae. albopictus* was predicted by a combination of macro- and microhabitat variables such as muddy water quality (*r*^2^=25.40) and developed medium intensity areas (*r*^2^=22.44) (Sallam et al. 2017b). These findings were consistent with other researches, which found a link between *Ae. albopictus* population density and water quality (e.g., organic muddy, dark turbid) (Higa et al. 2010, Nasir et al. 2017), developed medium intensity (Comiskey et al. 1999), and pH (Honório et al. 2006). Our findings were confirmed by the maximum *r*_2_ values generated by combined regression models (Tables 4 and 5). This finding broadly supports the work of other studies in this area linking microhabitat suitability with *Ae. albopictus* breeding in discarded tires (Comiskey et al. 1999, Vezzani et al. 2005, Maciel-de-Freitas et al. 2007, Vezzani and Albiocco 2009, Dória et al. 2010, Murrell et al. 2011). This mosquito vector was found to be affected by detritus and tires with high nutrient contents (Merritt et al. 1992, Mazine et al. 1996).

In contrast, the microhabitat factors predicted the presence of *Cx. quinquefasciatus*. In particular, although macrohabitat factors demonstrated high prediction gain (*r*^2^=6.0, AICc=1673), the combination of macro- and microhabitat variables improved the prediction gain, and the combined regression model was selected as the best model in delineating habitat factors for *Cx. quinquefasciatus* (*r*^2^=18.67, AICc=1566). This model indicated an intact/undamaged tire status and muddy/clear water quality as the best predictors for *Cx. quinquefasciatus*. *Culex. quinquefasciatus* also occurred at much lower densities and exhibited a much smaller geographic range that was indicated by the standard deviational ellipsoid. This finding supports evidence from the previous observations (Merritt et al. 1992, Qualls and Mullen 2006). However, the predicting variables of *Cx. quinquefasciatus* in water-containing habitats seemed to be context dependent in terms of interspecific comptetion with other mosquito populations, host preference, type of breeding sites and season (Sallam et al. 2017a). This mosquito species showed resilience to ecological requirements for their presence and abundance and demonstrated biological fitness to breed in a wide range of breeding sites. Another study need to be conducted to quantify all potential breeding sites of *Cx. quinquefasciatus* in the City of New Orelans to understand their predicting factors in their breeding sites.

Generally, the presence of both *Ae. albopictus* and *Cx. quinquefasciatus* in discarded tires was influenced much more by microhabitat factors than macrohabitat factors. This is important because mosquito dietetic necessities are met through the intake of both dead and living organic material (Merritt et al. 1992, Mazine et al. 1996, Yee et al. 2004). This finding warrants the need to not only reinforce discarded tire pick-up initiatives in New Orleans but also control of mosquito breeding in this urban landscape by exploring water quality conditions of *Aedes* and *Culex* larval habitats that proliferate mosquitoes. Our study is a preliminary step for a comprehensive model that integrates crowd-sourcing data with active surveillance to more completely understand Spatio-temporal distribution of container breeding mosquitos in their potential colonization sites.

## Conclusion

Our results indicate that although the problem of discarded vehicle tires is widespread (in both frequency and number) in the 13 planning districts of New Orleans, their center of concentration was found to be in Mid-city. We identified 13 mosquito species from 65 sites (38.46%) representing 1,136 (∼37.08%) sampled tires in seven of the planning districts. The two most dominant mosquito species, *Ae. albopictus* and *Cx. quinquefasciatus*, were found to co-exist in 21 sampling sites (∼32.31%) representing 214 tires (35.79%) from four planning districts: Bywater, Gentilly, Lower Ninth Ward, and New Orleans East. The clustered and random spatial distribution polygons of *Ae. albopictus* and *Cx. quinquefasciatus* were found to be in seven planning districts. Both species were significantly predicted by microhabitat characteristics rather than macrohabitats, especially *Ae. albopictus*. Given the diverse mosquito fauna of Louisiana, we recommend rigorous ecological studies be conducted to identify ecological and socioeconomic factors that may influence species composition within and among sites and their temporal distribution pattern in New Orleans. This is a critical step towards understanding the contribution of macro- and microhabitat characteristics in predicting colonized mosquito populations in water-holding tires. Also, this study adds to the growing body of research that indicates that different factors found in the tire environments influence oviposition and/or larval performance.

## Acknowledgments

We thank Matt Torri and the employees of the City of New Orleans Sanitation Department, the City of New Orleans 311 Call Center, and City of New Orleans Mosquito and Termite Control Board staff who helped to locate tire sites and assist with mosquito identification. A special thank you to all the property owners for allowing us access to study sites. This work was partially supported by Grant ZIKA-CDC-RFA-CK14-140199HF, building domestic surveillance, laboratory, vector control, and pregnancy registry capacity to respond to Zika virus. The views expressed in this article are those of the author and do not necessarily reflect the official policy or position of the Department of the Navy, Department of Defense, nor the U. S. Government.

## Data Availability

The authors declare that all the relevant data supporting the study findings are available from the City of New Orleans upon request.

## Authors’ contributions

TA, CSL, and CR carried out the experimental design and data sampling. MFS: carried out geo-database and model building, data analysis, and writing the first draft of the manuscript. IKM helped in validating models and editing manuscript. All authors approved the final version of the manuscript.

## Competing interests

The authors declare that they have no competing interests.

## Ethics approval and consent to participate

Not applicable

